# Genome-wide maps of distal gene regulatory regions active in the human placenta

**DOI:** 10.1101/364026

**Authors:** Joanna Zhang, Corinne N. Simonti, John A. Capra

## Abstract

Placental dysfunction is implicated in many pregnancy complications, including preeclampsia and preterm birth (PTB). While both these syndromes are influenced by environmental risk factors, they also have a substantial genetic component that is not well understood. Precisely controlled gene expression during development is crucial to proper placental function and often mediated through gene regulatory enhancers. However, we lack accurate maps of placental enhancer activity due to the challenges of assaying the placenta and the difficulty of comprehensively identifying enhancers. To address the gap in our knowledge of gene regulatory elements in the placenta, we used a two-step machine learning pipeline to synthesize existing functional genomics studies, transcription factor (TF) binding patterns, and evolutionary information to predict placental enhancers. The trained classifiers accurately distinguish enhancers from the genomic background and placental enhancers from enhancers active in other tissues. Genomic features collected from tissues and cell lines involved in pregnancy are the most predictive of placental regulatory activity. Applying the classifiers genome-wide enabled us to create a map of 33,010 predicted placental enhancers, including 4,562 high-confidence enhancer predictions. The genome-wide placental enhancers are significantly enriched nearby genes associated with placental development and birth disorders and for SNPs associated with gestational age. These genome-wide predicted placental enhancers provide candidate regions for further testing in vitro, will assist in guiding future studies of genetic associations with pregnancy phenotypes, and aid interpretation of potential mechanisms of action for variants found through genetic studies.

## INTRODUCTION

The placenta is a complex temporary organ, essential for successful pregnancy. The placenta performs many vital functions including transfer of nutrients to the developing fetus and protection against infectious agents [1]. Placental dysfunction has been connected to pregnancy complications, such as preeclampsia and preterm birth (PTB) [2–5]. PTB and preeclampsia both have environmental risk factors as well as a genetic component that is not well understood. Family and pedigree studies of PTB and preeclampsia suggest strong genetic components, but heritability estimates for both vary considerably [5,6], and genetic associations found through genome-wide association studies (GWAS) of these and other disorders of pregnancy have been difficult to regulate [7,8]. Though a recent study of more than 43,000 women has identified and replicated several loci associated with gestational duration and preterm birth [9].

Precisely controlled gene expression during pregnancy is crucial to proper development, and these gene regulatory “programs” are mediated by enhancers, gene regulatory elements that play a large role in development and thus disease [10–12]. Disruption of enhancers and gene regulation have been shown to influence risk for many complex diseases [10,13]. Thus, mapping the enhancer landscape is a common step in the search for and interpretation of genetic associations. As is common for complex diseases, the genetic variants that have been implicated in PTB risk by GWAS are non-coding and thus difficult to interpret. Typical enhancer identification methods are impractical in early placental stages for many reasons, but perhaps most importantly because sampling the placenta increases risk of pregnancy loss [14]. *In vivo* studies in model organisms have lent insight to early placental development, but the rapid evolution of pregnancy across taxa often limits the translatability of this work [15].

To address the challenge of mapping gene regulatory elements active in the placenta, we used the EnhancerFinder [16] machine learning approach to predict placental enhancers. Using computational methods to synthesize existing functional studies, transcription factor (TF) binding, and evolutionary information to identify enhancers avoids many of the difficulties of studying the placenta discussed above. Indeed, such methods have historically been successful in identifying and interpreting regulatory regions [16–18]. We present a set of 4,562 placental enhancers predicted genome-wide. These putative enhancers show clear relevance to placental biology; they are located near many genes involved in placental function and development and are significantly enriched for genetic variants associated with pregnancy phenotypes and complications. These predicted enhancers provide candidate regions for researchers to test *in vitro*, and propose mechanisms of action for variants found through GWAS. To facilitate their use, all the enhancer predictions are integrated into GEneSTATION (v2.0) [19].

## RESULTS

### A two-step machine-learning framework for placental enhancer prediction

To predict placental enhancers, we used the EnhancerFinder algorithm, which integrates sequence, evolutionary, and functional properties of known enhancers to build statistical models that enable the identification of new enhancers [16]. This approach proceeds in two steps. First, a model is built to distinguish known enhancers active in any cellular context from regions from the genomic background (Step 1). Then, models for classifying enhancers active in particular tissues are trained by comparing enhancers active in a tissue of interest to enhancers only active in other tissues (Step 2). This two-step approach yields more specific predictions than a single step approach [16].

We trained our classifiers using enhancers defined by cap analysis of gene expression (CAGE) from the FANTOM5 Transcribed Enhancer Atlas [20]. Analyzing 411 different tissues and cell lines, they identified 38,538 robust human enhancers, of which 748 were active in the human placenta. We characterized each enhancer by its DNA sequence properties, evolutionary conservation, and chromatin state. Each region’s DNA sequence composition was quantified by counting the occurrence of all five-nucleotide-long (5-mer) DNA sequences within the region. Evolutionary conservation was quantified using mammalian conserved elements from phastCons [21]. Finally, we used functional genomics data from the Roadmap Epigenomics Project [22], including histone modifications and DNaseI hypersensitivity data from hundreds of cellular contexts, to quantify the chromatin state of the region. (See the Methods for a complete description of the features.).

Then, using these features, we trained a multi-kernel support vector machine (SVM) classifier—with one kernel for each of the three data types—to distinguish robust enhancers from random, length-matched non-enhancer regions from the genomic background (Fig 1; Step 1). For Step 2, we trained a placental enhancer classifier using the 748 known placental enhancers as positives and a random subset of 2,000 robust non-placental enhancers as the negatives (Fig 1).

**Fig 1.**
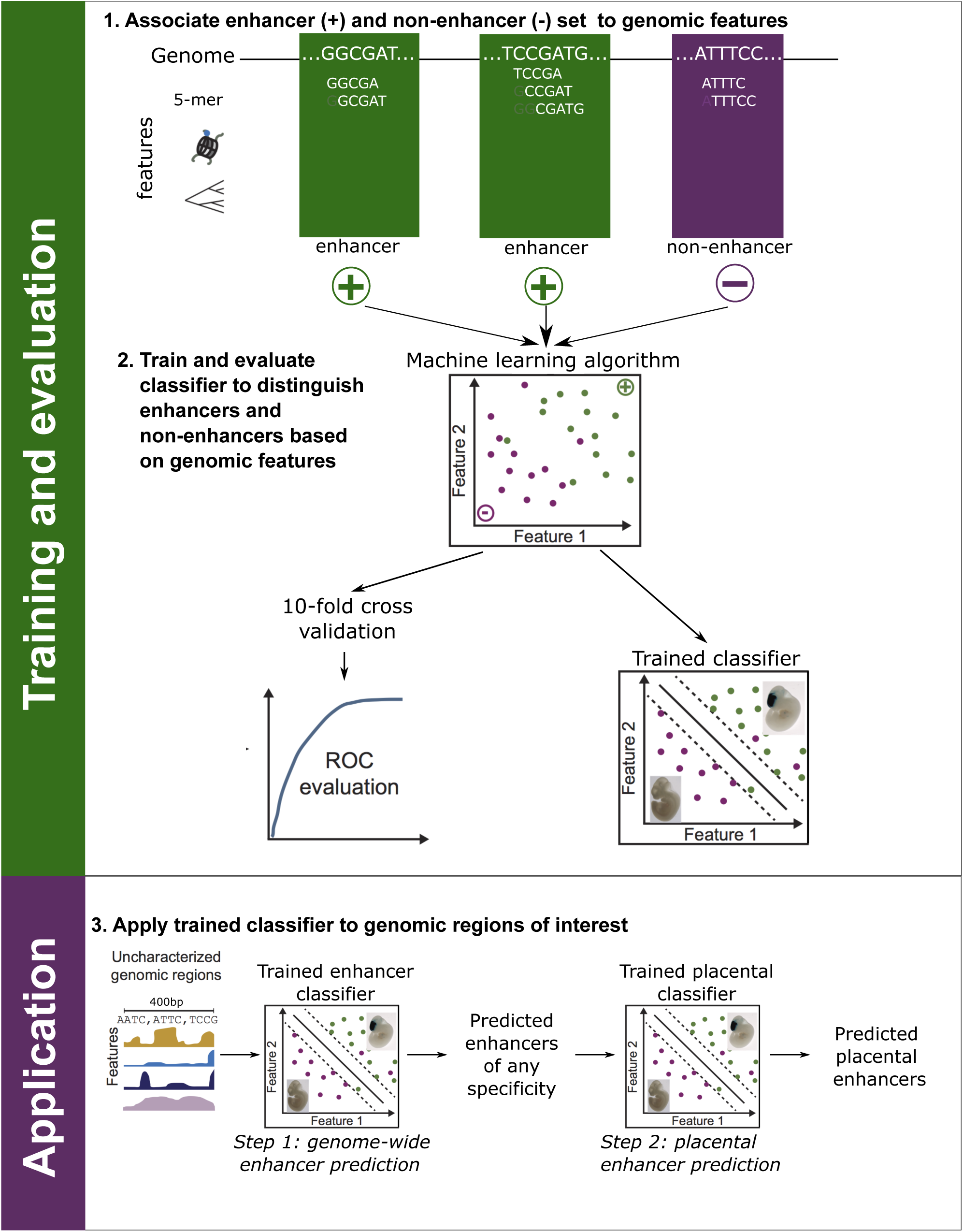
Schematic of the placental enhancer prediction pipeline. First, we associated known enhancers from diverse tissues (+) and non-enhancer regions from the genomic background (–) with a range of informative features including their DNA sequence patterns, functional genomics data, and evolutionary conservation across species. Second, we trained a multi-kernel support vector machine to distinguish the enhancers from regions without enhancer activity using the associated features. We evaluated the performance of trained classifiers using 10-fold cross validation. Finally, we applied a classifier trained to distinguish enhancers from non-enhancers to all sequences in the human genome (Step 1). Then we applied a second classifier trained to distinguish placental enhancers from enhancers active in other tissues (Step 2). This produced an accurate set of genome-wide placental enhancer predictions.

### Accurate prediction of known placental enhancers

To assess the performance of our trained classifiers, we used 10-fold cross validation to compute average receiver operating characteristic (ROC) curve and precision-recall (PR) curves. In 10-fold cross validation, ten models are trained using a different 90% of the positive and negative training regions, and then each model is evaluated on remaining 10% of the regions. We quantified our method’s overall performance by the average area under the curve (AUC) over the 10 runs.

The trained Step 1 classifier performs very well at identifying FANTOM enhancers from genomic background (Fig 2A; ROC AUC=0.93, PR AUC=0.78). The classifier trained to distinguish placental enhancers from enhancers active in other contexts (Step 2) also has strong performance (Fig 2B; ROC AUC=0.84, PR AUC=0.70). While distinguishing enhancers active in the placenta from enhancers active in other tissues is more challenging than generally distinguishing enhancers from the genomic background, our approach still performs well at this task.

**Fig 2.**
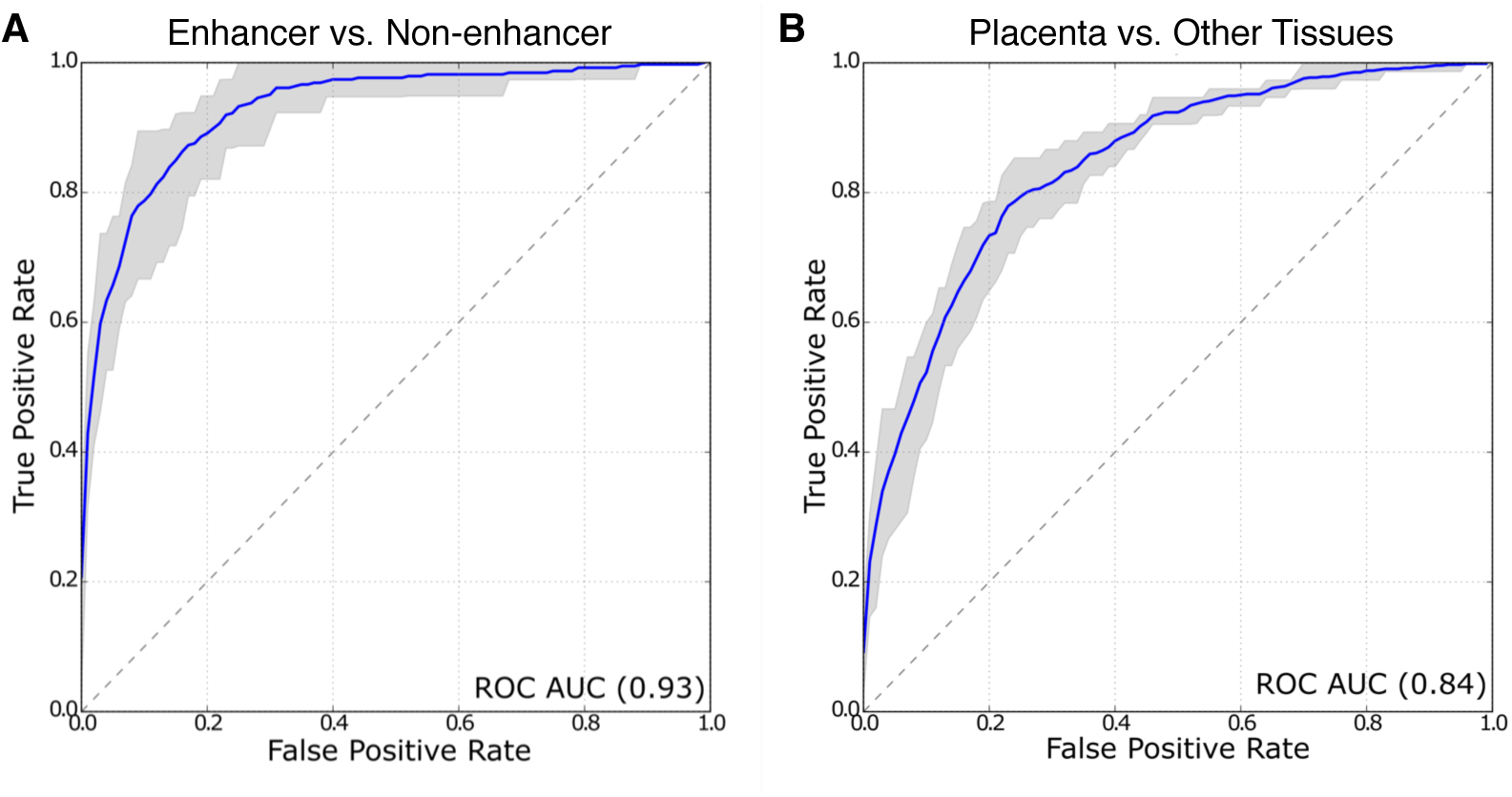
The trained classifiers accurately identify placental enhancers. Receiver operating characteristic (ROC) curves for the classifiers trained to distinguish enhancers from non-enhancers (A, Step 1) and placental enhancers from enhancers active in other tissues (B, Step 2). Both perform significantly better than expected by chance with areas under the ROC curve (AUC) of 0.93 and 0.84 respectively. The shaded region represents the performance range observed over the 10 cross validation runs. The diagonal line represents chance performance. The corresponding Precision-Recall curve AUCs are 0.78 and 0.70, respectively.

### Functional genomics data from pregnancy-related tissues are the most informative for distinguishing placental enhancers from other enhancers

To investigate the genomic attributes most useful to the placental enhancer classifier, we examined the individual feature weights the algorithm assigned in the functional genomics kernel after Step 2 training. A positive feature weight indicates association with placental enhancer activity, while a negative feature weight is associated with enhancer activity in another context. The most informative contexts (i.e., the contexts whose histone modification features had the largest absolute weights) within the kernel were from placental and related tissues (trophoblast cells, amnion, and endometrial stromal cells), and the least informative features came from cellular contexts unrelated to pregnancy (Fig 3).

**Fig 3.**
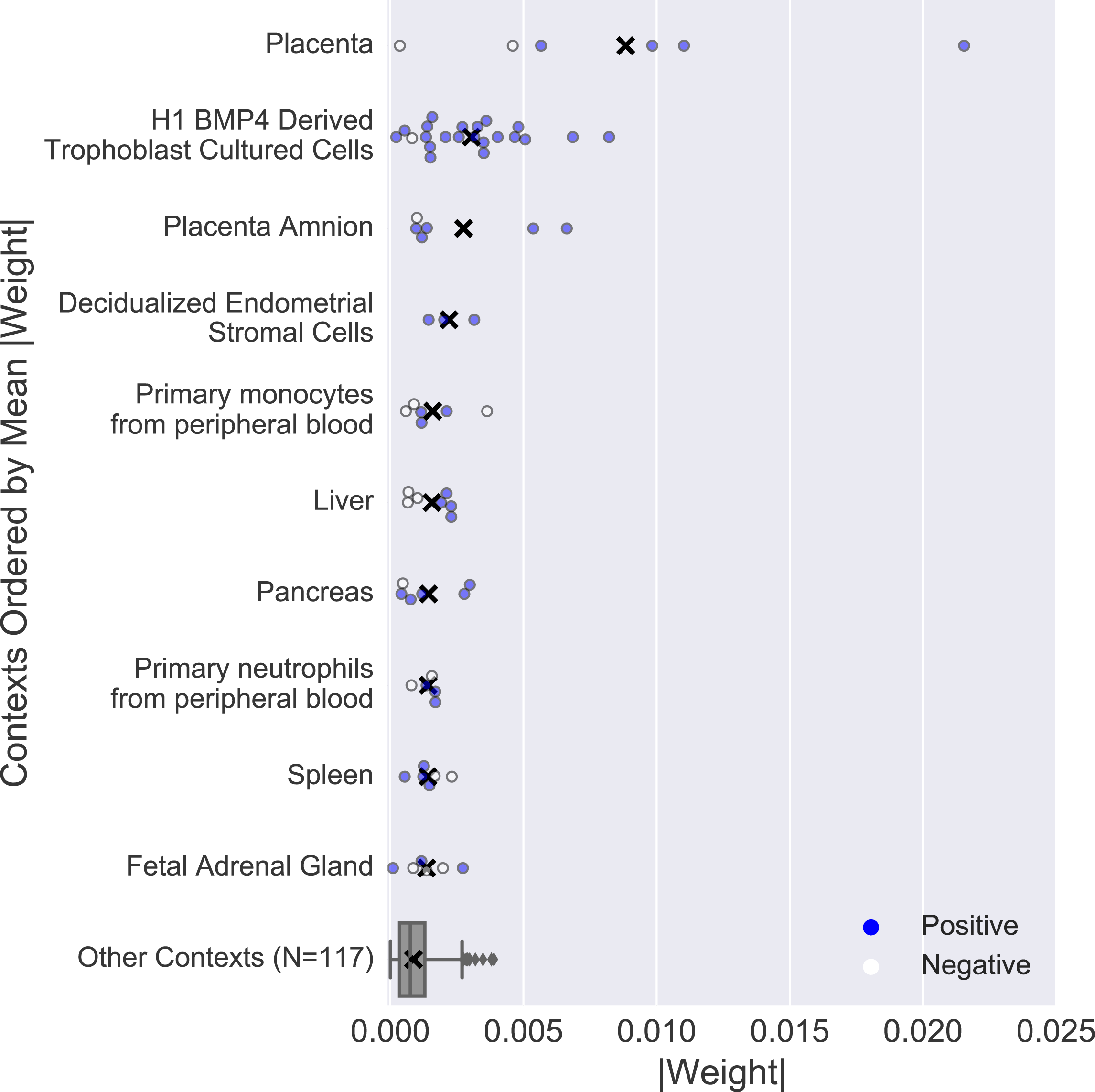
Functional genomics data from pregnancy-related tissues are highly weighted by the placental enhancer classifier. The absolute value of the weight assigned to each functional genomics feature in the SVM is plotted (positive weight: blue, negative weight: white, mean of absolute weights: black X). The absolute weights on the functional genomics features from the other 117 contexts were collapsed into one box plot (outliers are plotted as gray diamonds).

### A genome-wide map of regions with potential placental regulatory activity

To identify genomic regions with potential placental regulatory activity genome-wide, we applied our trained classifiers to the human genome by tiling all human chromosomes into regions the length of an average FANTOM5 placental enhancer (400 bp) overlapping by 200 bp. We filtered out tiles that overlapped gaps in the genome assembly, exons, and likely promoter regions (5 kb region upstream of each transcription start site). Tiles assigned to both the enhancer and placental enhancer by the SVM classifiers were considered putative placental enhancer. Those with strong predictions in both classifiers (SVM score > 1) were considered high confidence putative placental enhancers. Merging overlapping tiles yielded 4,562 high-confidence placental enhancers, covering 3,475,438 bp of the genome, and 33,010 putative enhancers, covering 38,893,990 bp of the genome (Table 1).

**Table 1.** Statistical summary of genome-wide placental enhancer predictions.

| Enhancer set | Count | Mean length (bp) | Genome Coverage (bp) |
| --- | --- | --- | --- |
| High Confidence Placental Enhancers | 4,562 | 762 | 3,475,438 |
| Potential Placental Enhancers | 33,010 | 846 | 38,893,990 |

### Predicted placental enhancers are enriched near genes with placental functions

To evaluate the relevance of our high-confidence predicted placental enhancers to placental biology and pregnancy, we examined nearby genes in the context of known gene annotations. Using the functional enrichment analysis tool GREAT [23], we mapped each region to putative gene targets and then tested for the enrichment of relevant Gene Ontology (GO) functional annotations. We found significant enrichment for many relevant terms such as “placenta development” and “decreased placental labyrinth size” (selected terms: Table 2, full list: Supplementary Table 1).

**Table 2.** Placenta-relevant functions significantly enriched among genes near high-confidence predicted placental enhancers. GO BP = Gene Ontology Biological Process.

| Ontology | Term | Binomial Fold Enrichment | Binomial FDR Q-value |
| --- | --- | --- | --- |
| GO BP | Placenta development | 2.0 | 6.6e−13 |
| GO BP | Embryonic placenta development | 2.2 | 1.0e−12 |
| Mouse Phenotype | Decreased placental labyrinth size | 4.8 | 2.9e−33 |
| Mouse Phenotype | Abnormal placenta labyrinth morphology | 2.4 | 1.5e−28 |
| MGI Expression | TS4 Zona Pellucida | 2.1 | 3.9e−64 |
| Disease Ontology | Neoplasm of body of uterus | 2.7 | 3.5e−24 |
| Disease Ontology | Persistent fetal circulation syndrome | 4.8 | 1.7e−06 |
| Disease Ontology | Newborn respiratory distress syndrome | 2.6 | 3.2e−06 |

### Predicted placental enhancers are enriched for regions associated with gestational age and preterm birth

To assess the biological importance of our high-confidence placental enhancers, we tested for enrichment of regions associated with gestational age and preterm birth in a recent genome-wide association study (GWAS) [9]. Forty-three of our predicted enhancers overlapped 12 out of 14 GWAS regions. To interpret this, we compared the observed overlap to the number of overlaps found for 10,000 randomly generated sets of genomic regions length- and chromosome-matched to our predictions and excluding genomic gaps. Our putative enhancers were significantly enriched for relevant GWAS catalogued regions associated with preterm birth and gestational age (*P* < 0.0001) with a calculated fold enrichment of 2.69 (relative to the mean of the randomized sets).

To compare the high-confidence placental enhancer set to the candidate placental enhancer set, we tested the enrichment for specific functions near the candidate regions using GREAT and for overlap with the pregnancy-related GWAS regions. We found similar placenta-related GO terms enriched near the larger candidate placental enhancer set, for example: with GO terms such as “placenta development” (*P* = 3.80e–147) and “embryonic placenta development” (*P* = 3.82e–99). The candidate enhancers were also enriched for GWAS regions associated with preterm birth and gestational age (relative fold enrichment: 2.23, *P* < 0.0001). Thus, there is evidence to suggest that additional regulatory regions relevant to placental biology are present in the candidate set.

### Predicted placental enhancers expand previously published placental enhancer datasets

We further compared our placental enhancer predictions to a recently published set of 2,216 computationally predicted placental enhancers [17]. These candidates were identified by identifying TFs implicated in placental and trophoblast function by GREAT and then predicting enhancer activity based on clustering of TF binding sites (TFBS) in the mouse genome. We will refer to these putative enhancers as “TFBS clusters.”

We calculated the overlap between the TFBS clusters that mapped to human genome and did not overlap exons or a 5kb region upstream of TSSs (1,044 TFBS clusters) and our high-confidence placental enhancers. We found 82 elements (20,154 bp) overlapped between the two sets. Because the biological information used to define enhancers differed between the sets, it is not surprising that our predictions and the TFBS clusters identify largely distinct regions of the genome.

To evaluate the functional relevance of the TFBS clusters, we tested for enriched relevant functions using GREAT and for enrichment in overlap with preterm birth and gestational age GWAS regions. We examined the GO biological process terms “placenta development” and “embryonic placental development” and both were comparably enriched among genes near the TFBS clusters (*P* = 2.95e–15 and *P* = 2.33e–17, respectively) as among our predicted enhancers. The results were similar for enrichment for pregnancy-related GWAS regions. While 43 of our placental enhancers fell within a GWAS region associated with preterm birth and gestational age with a calculated fold enrichment of 2.69 (*P* < 0.0001), the TFBS clusters overlapped 13 elements had a fold enrichment of 3.07 (*P* < 0.0006). Overall, comparing the significant functional annotations of the TFBS clusters with our predicted placental enhancers revealed similar levels of enrichment for relevant functional terms.

### Placental enhancers are enriched for ancient transposable elements

Transposable elements (TEs) often create regulatory elements in pregnancy-related tissues [24–26]. We calculated the enrichment of the FANTOM placental enhancers as well as both predicted sets for overlap with TEs. Overall, as expected due to the silencing of TEs across the genome, each set is significantly depleted of TEs (*P* < 0.001, randomization test) compared to the genomic expectation. However, the age distribution of TEs present in the placental enhancers compared to TEs overlapped by permuted enhancer sets is significantly enriched for TEs originating in the common ancestor of theria or before (Fig 5; *P* < 0.001, randomization test). The enrichment for ancient TEs and depletion of more recent TEs is a common pattern across validated enhancers [27], and thus the similar observation across our predicted enhancers lends support to their enhancer activity.

**Fig 4.**
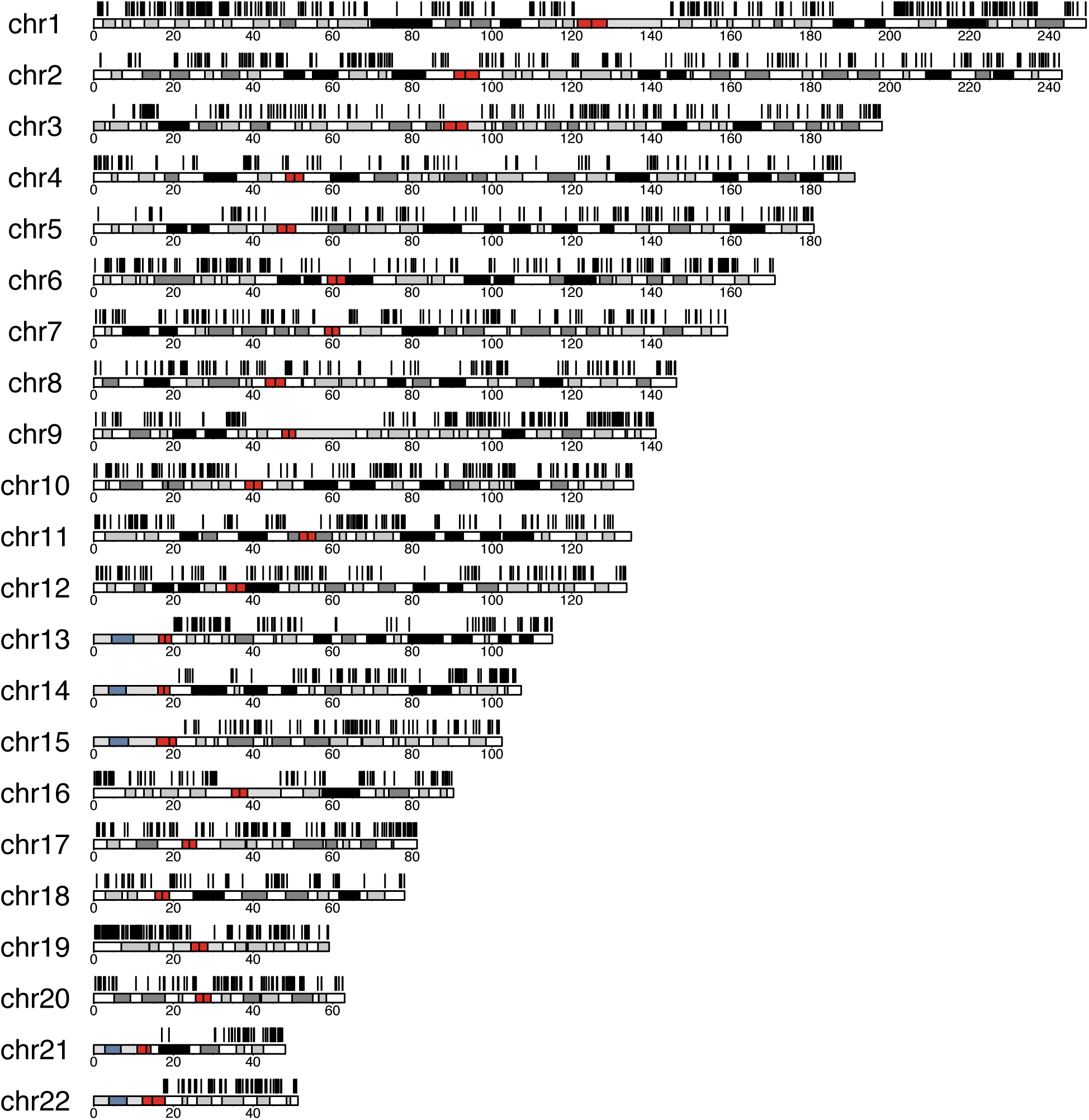
High-confidence predicted placental enhancers are found across the human genome. The black lines indicate the locations of a high-confidence predicted placental enhancer on the human chromosomes. We predicted 4,562 high confidence placental enhancers and 33,010 potential placental enhancers (Supplementary Files 1 and 2).

**Fig 5:**
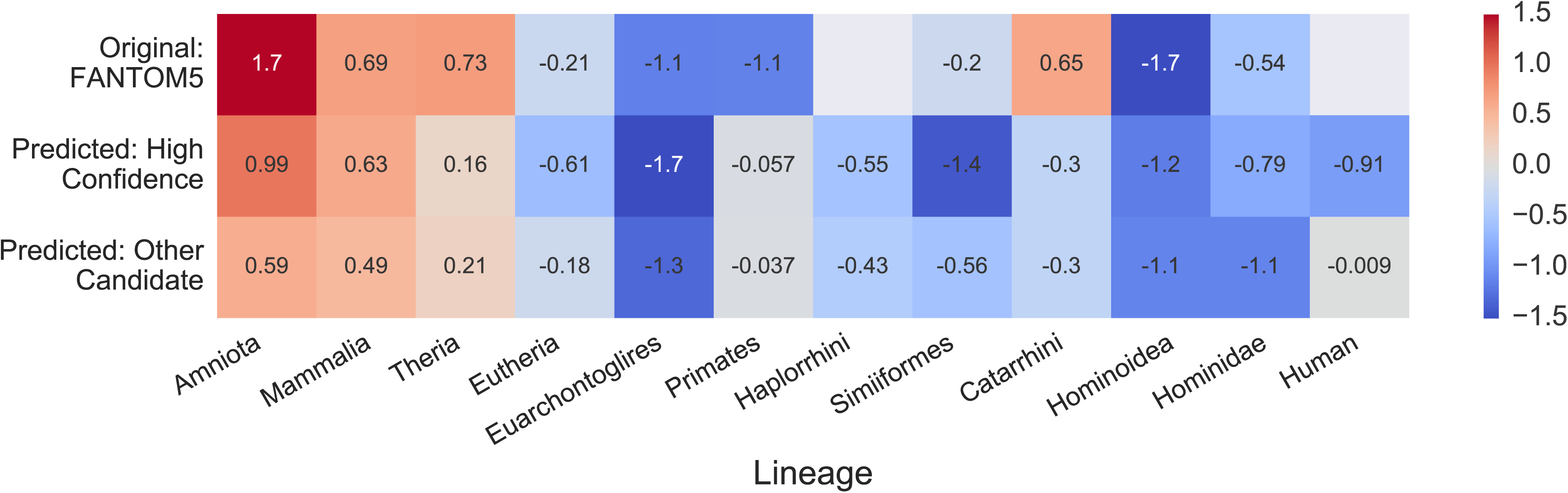
Validated and predicted placental enhancers are enriched for ancient transposable elements. We computed the enrichment for overlap of transposable elements (TEs) with origins on different lineages for experimentally validated and predicted enhancer sets. The enrichment was computed in reference to the mean of the genome-wide overlap observed in 1,000 (predicted) or 10,000 (FANTOM5) permuted enhancer sets. The log_2_ of the relative change is given for each comparison. Asterisks indicate significant enrichment (*P* < 0.05, randomization test). Empty gray boxes indicate there were not enough enhancers to test for enrichment.

## DISCUSSION

Using an established machine learning framework, we identified 4,562 high-confidence placental enhancers, as well as an expanded set of 33,010 candidate placental enhancers. These putative regulatory regions are enriched near genes relevant to pregnancy, are enriched for overlap with variants associated with diseases of pregnancy, and have similar transposable element profiles as validated enhancers. In addition, the predicted enhancers significantly expand previously published sets of placental enhancers, and thus provide greater power to interpret genetic associations with diseases influenced by the placenta. For example, the fact that 12 out of 14 regions associated pregnancy complications in a recent GWAS are in high linkage disequilibrium with a predicted enhancer underscores the utility of these genome-wide enhancer maps. These candidates suggest targeted regions for testing when seeking the causal variants in these regions and dissecting how they influence pregnancy. More accurate interpretation of these and future GWAS hits is necessary for understanding the complex biology of pregnancy and eventually improving the identification and prevention of disorders such as preterm birth. To facilitate the use of our enhancer maps, they are now integrated into the GEneSTATION web platform for studying pregnancy and preterm birth [19].

Our predicted enhancer maps can be improved in several dimensions. First, they are undoubtedly incomplete. Enhancer activity is highly context and stimulus dependent. Due to the paucity of training data from diverse contexts, we have focused on identifying a set of candidate regions that have hallmarks of potential regulatory activity in the placenta broadly without making specific contextual predictions. Furthermore, the patterns learned by our machine learning classifier generalize existing patterns in the evolution, sequence, and functional genomics of known placental enhancers, but are constrained by what is currently known. Finally, there is heterogeneity in the cellular makeup of the placenta and existing data do not enable cell-specific predictions. As more enhancer data become available from relevant cellular contexts, we will continue to refine our predictions and integrate them with other annotations.

While the costs and technical difficulties of agnostically identifying enhancers are decreasing, many tissues and cell types remain difficult to assay due to biological constraints and ethical considerations. These challenges are compounded for tissues like the placenta that are rapidly evolving between species, limiting the utility of information garnered through the study of model organisms. Computational approaches, such as those presented here, paired with growing collections of experimentally validated regulatory regions provide a promising avenue for enabling researchers to interrogate the gene regulatory architecture of the placenta and other tissues that are difficult to assay.

## METHODS

### Genome-wide placental enhancer predictions

We based our approach on the EnhancerFinder two-step machine learning algorithm for predicting enhancers and their tissues of activity. We first trained an SVM classifier based on diverse sequence, evolutionary, and functional genomics features to distinguish known enhancers active in a range of tissues from the genomic background. Then in the second step, additional classifiers were trained to distinguish enhancers active in different tissues from one another. In this step, all enhancers active in a tissue of interest (placenta) are used as positive training examples and all enhancers not active in the tissue are treated as negatives.

#### Training regions

We downloaded the hg19 genomic locations of all 38,538 robust human enhancers identified by CAGE from the FANTOM5 Transcribed Enhancer Atlas. The data included 748 human placental enhancers. The average length of a FANTOM5 placental enhancer is 400 bp.

To train the enhancer classifier (step 1), the positive set consisted of a random subset of 385 robust human enhancers (fixed to a length of 400 bp at the center of any enhancer). Our negative set consisted of 2,000 random genomic regions matched to the length and chromosome distribution of the positive set and excluding FANTOM5 enhancers and hg19 genome assembly gaps. The random genomic regions were generated using shuffleBed [28]. To train the placental enhancer classifier (step 2), we used the 748 human placental enhancers (fixed at a length of 400 bp from each enhancer center) as positives. The negative set consisted of a random subset of 2,000 robust human enhancers, excluding placental enhancers. All analyses in this paper were performed in reference to the UCSC Genome Browser February 2009 assembly of the human genome (GRCh37/hg19). Any dataset not in this build was mapped over to hg19 coordinates using the liftOver tool from the UCSC Kent tools with default parameters [29].

#### Feature data

Three types of data were used as features in the MKL algorithm: functional genomics, evolutionary conservation, and DNA sequence motifs. Each type of data was assigned to its own kernel. Following the approach used in previous applications of EnhancerFinder [16], we used linear kernels, consisting of computed dot products of feature vectors, for the functional genomics and evolutionary conservation data. For the DNA sequence-based features we used a 5-spectrum kernel. The MKL algorithm combines the three kernels by learning weights to assign to each kernel from the training set [16].

For the functional genomics kernel, we obtained 980 histone modification datasets (H3K27ac, H3K4me1, H3K4me4, etc.) and 39 DNase datasets from 128 cellular contexts in the Human Epigenome Atlas [22], as well as H3K27ac, H3K4me3, and DNaseI peaks identified in decidualized endometrial stromal cells from Lynch *et al* [24]. Feature vectors were constructed by overlapping genomic regions in the training set with each functional genomics dataset. Each region was associated with a binary vector that represented the presence or absence of overlap with each feature dataset. We took evolutionary conservation scores from the UCSC Genome Browser phastConsElements46way tracks for placental mammals, primates, and vertebrates. Each genomic region was assigned the highest conservation score of any overlapping phastCons element. Genomic regions not overlapping a phastCons element were assigned a score of zero. To quantify the DNA sequence of a region of interest, we counted the occurrence of all possible length 5 bp DNA sequence motifs (5-mers) within genomic regions of interest.

#### Classifier training and prediction

All classifiers were trained using the Multiple Kernel Learning (MKL) functionalities of the SHOGUN Machine Learning Toolbox [30]. The algorithm uses features of the training set to learn a linear function that separates positives from negatives. Genomic regions can then be assigned a score based on their position relative to the separating hyperplane learned by the SVM. A positive score indicates that the region belongs to the positive set, while a negative score indicates membership in the negative set. The magnitude of the score indicates the confidence the algorithm places on its prediction. Only regions that are predicted to be positives by both classifiers are considered candidate placental enhancers.

#### Classifier evaluation

We evaluated the performance of our trained classifiers using 10-fold cross validation and computing ROC curves and precision-recall (PR) curves averaged over folds. In a 10-fold cross validation, the training data are partitioned into 10 equal subsets, and the classifier is trained 10 times. Each time, only 9 of the 10 subsets are used to train the classifier. The trained classifier is then applied to the held-out subset and evaluated based on the true status of these regions. The performance of the classifier is then quantified using ROC AUC and a PR AUC.

#### Interpreting Algorithm Weights for the Functional Genomics Kernel

Based on positive and negative training data, our algorithm reports the kernel and feature weights learned during training. The total kernel weight is computed along with the weight for each individual feature weight within that kernel. Positive values are assigned to features associated with the positive input set and features associated with the negative input set score more negatively. After training our placental enhancer classifier (Step 2), we examined the individual weights within its functional genomics kernel to determine whether placenta-related histone modifications were weighted higher than histone modifications found in other cellular contexts. In this case, positive weights are associated with placental enhancer activity and negative weights are associated with enhancer activity in other cellular contexts.

#### Genome-wide Placental Enhancer Prediction

To predict placental enhancers genome-wide, we tiled each autosome into 400 bp regions (the average length of a FANTOM placental enhancer) in overlapping increments of 200 bp. We omitted the sex chromosomes from our analyses. These regions were filtered to remove any tiles that overlapped an exon or fell within 5 kb of a transcription start site (TSS) to minimize association with promoter regions. Coordinates for exons and TSSs were downloaded from the Ensembl GRCh37 Feb 2014 [31] using the Biomart archive. We applied the trained enhancer and placental enhancer classifiers to all remaining tiles. We merged all overlapping regions that received scores greater than zero from both the enhancer and placental enhancer classifiers. The resulting 33,010 merged regions are our candidate placental enhancer set. To obtain a refined list of predicted regions, we fixed a minimum threshold score of greater than one from both of our trained classifiers. After merging overlapping regions that met our criteria, a subset of 4,562 candidate placental enhancers remained and became our high-confidence placental enhancer set.

### Analysis of genome-wide placental enhancer predictions

#### Gene ontology annotation enrichment

To identify the functional annotations, phenotypes, and pathways enriched among genes nearby the predicted placental enhancers, we used GREAT with the default settings. GREAT is a web tool that takes a set of genomic regions and associates them with their putative target genes and target gene annotations [23]. GREAT calculates the enrichment of annotations within the input regions and returns the terms that are significantly enriched near the input regions. We submitted our candidate placental enhancer set as well as our high confidence placental enhancer set to GREAT, using the default entire human genome as the background.

#### Enrichment for regions relevant to pregnancy

We calculated the enrichment for GWAS SNPs in our candidate placental enhancer set and high-confidence placental enhancer set. We obtained 14 preterm birth and gestational age GWAS regions (omitting 3 regions on the X chromosome) from a recent GWAS [9]. For each set of enrichment analyses, we generated 10,000 sets of random genomic regions that were matched to the predicted enhancer set based on the length and chromosome distribution. Then, we computed the overlap of each of the 10,000 random region sets with each set of regions of interest. Enrichment was calculated by dividing the overlap of our predicted set with the mean overlap of the 10,000 randomly generated sets, and an empirical p-value was obtained by counting the number of random sets for which as much or more overlap with the regions of interest is observed.

#### Comparison to previous placental enhancer predictions

We downloaded a set of 2,216 placental enhancers defined using transcription factor binding site (TFBS) clusters related to placental function from supplementary material of Tuteja et. al [17]. Of the 2,216 TFBS clusters whose build was of the UCSC Genome Browser July 2007 assembly of the mouse genome (NCBI37/mm9), 2,207 TFBS clusters mapped into hg19 using liftOver [29]. From these TFBS clusters, we generated a subset of 1,044 regions by filtering out regions overlapping exons and regions within 5 kb of a transcription start site (TSS). The motivation for generating a smaller subset of TFBS clusters comes from our concern that predicted placental enhancers defined by TFBSs nearby TSSs may have an increased chance of being associated with promoters rather than enhancers. All enrichment tests were calculated on both the larger and smaller subset of TFBS clusters. Both sets of TFBS clusters had comparable enrichments. We report them for the smaller set that is more comparable to our enhancer sets here.

#### Transposable element enrichment analysis

TE genomic locations were retrieved from RepeatMasker v4.0.5 [32]. The clades in which each TE is present were taken from Dfam v1.4 [33]. In situations where Dfam provided multiple clades, the clade of the most recent common ancestor was designated as the origin. We collapsed all TEs originating in the last common ancestor of amniota or before into one category.

For both the FANTOM5 placental enhancers and the high-confidence predicted placental enhancers, we used shuffleBed [28] to shuffle enhancer regions around the genome. We constrained the shuffled regions to the chromosome of the corresponding observed region and did not allow shuffled regions overlap one another, gaps in the genome assembly, or ENCODE blacklist regions [34]. For the FANTOM5 enhancers, we created 10,000 sets of shuffled regions. For the predicted enhancers, we created 1,000 sets of shuffled regions separately for the high-confidence and candidate sets. We calculated the permutation-based p-value for each lineage of origin for all TEs by calculating the number of permuted sets that overlapped more or the same amount of TEs appearing on a given lineage. Tests were only performed if at least 10 enhancers overlapped a TE of the given lineage.

## ACKNOWLEDGMENTS

We thank members of the Capra and Rokas Labs for helpful discussions. We thank Ge Zhang for sharing information on the gestational age and preterm birth GWAS. This work was supported by NIH grants R01GM115836 and R35GM127087 to JAC, an Innovation Catalyst Award from the March of Dimes Ohio Collaborative to JAC, and a Burroughs Wellcome Fund Preterm Birth Initiative Award to JAC. JZ was supported by a Vanderbilt Undergraduate Summer Research Program (VUSRP) Fellowship.

